# A comparative analysis of stably expressed genes across diverse angiosperms exposes flexibility in underlying promoter architecture

**DOI:** 10.1101/2023.06.12.544596

**Authors:** Eric J.Y. Yang, Cassandra J. Maranas, Jennifer L. Nemhauser

**Affiliations:** University of Washington, Department of Biology, Seattle, WA 98105-1800, USA

## Abstract

Promoters regulate both the amplitude and pattern of gene expression—key factors needed for optimization of many synthetic biology applications. Previous work in *Arabidopsis* found that promoters that contain a TATA-box element tend to be expressed only under specific conditions or in particular tissues, while promoters which lack any known promoter elements, thus designated as Coreless, tend to be expressed more ubiquitously. To test whether this trend represents a conserved promoter design rule, we identified stably expressed genes across multiple angiosperm species using publicly available RNA-seq data. Comparisons between core promoter architectures and gene expression stability revealed differences in core promoter usage in monocots and eudicots. Furthermore, when tracing the evolution of a given promoter across species, we found that core promoter type was not a strong predictor of expression stability. Our analysis suggests that core promoter types are correlative rather than causative in promoter expression patterns and highlights the challenges in finding or building constitutive promoters that will work across diverse plant species.

## Introduction

Precise control over gene expression is essential for development and survival. One of the first regulatory steps in expression regulation is transcription initiation, which is controlled by DNA regions designated as promoters. Current understanding of eukaryotic promoters is still remarkably limited, and we have difficulty even identifying a precise promoter region given an arbitrary sequence (Donczew & Hahn, 2017). A core promoter region is functionally defined as the minimal region required for transcription initiation, associated with binding of RNA Polymerase II (RNAPII) and General Transcription Factors (GTFs). Proximal and distal cis-regulatory elements contribute to the modulation of the core promoter’s activity and give it its characteristic expression profile. A sequence containing the proximal cis-regulatory elements as well as the core promoters is often referred to as the “promoter” region (Andersson & Sandelin, 2020; Biłas et al., 2016; Haberle & Stark, 2018; Schmitz et al., 2022). In practice, cloning and analysis projects often pick an arbitrary length (e.g., up to 2000 base pairs or until the next coding sequence) upstream of the transcription start site to define as the promoter region (Andersson & Sandelin, 2020; Schmitz et al., 2022).

Many core promoter elements have been identified within the core promoter region that are important in directing RNAPII and determining the transcription start site (TSS). The TATA-box motif is the most well-understood of the core promoter elements, yet TATA-box-containing promoters only account for about 20% of eukaryotic promoters and about 30% of *Arabidopsis* promoters (Donczew & Hahn, 2017; Molina & Grotewold, 2005). In plants, additional core promoter types were proposed by Yamamoto and colleagues based on their identification of over-represented motifs around a fixed distance from the transcription start site (Yamamoto et al., 2007, 2009). Y patch, or pyrimidine patch, motifs are C and T rich motifs whose presence had been recently shown experimentally to associate with stronger expression (Jores et al., 2021). CA and GA are additional core promoter elements, represented in approximately 20% and 1% of genic promoters, respectively (Yamamoto et al., 2009). Unlike the TATA-box which has a known GTF-binding protein associated with it, the molecular mechanism of the Y patch, CA and GA elements remain largely unknown. Core promoters that do not contain any of the identified core promoter types have been termed Coreless (Yamamoto et al., 2009, 2011). In *Arabidopsis*, Coreless promoters tend to be expressed more weakly but more broadly than those that contain TATA-boxes (Das & Bansal, 2019; Yamamoto et al., 2011).

Constitutive promoters, defined here as promoters that are on in all tissues at all times, are versatile tools in synthetic biology due to their desirable expression pattern (Yang & Nemhauser, 2022; Zhou et al., 2023). They are often used to drive expression of components used in synthetic circuits or metabolic engineering (Brophy et al., 2022; Patron, 2020; South et al., 2019; Wu et al., 2014). Core promoter regions of constitutive promoters (such as the Cauliflower Mosaic Virus 35S promoter) have often been used as the starting point to build synthetic promoters by introducing natural cis-elements or synthetic TF-binding sites upstream of these core promoter regions to artificially tune expression strength or confer new expression patterns (Ali & Kim, 2019; Belcher et al., 2020; Brophy et al., 2022; Brückner et al., 2015; Cai et al., 2020; Moreno-Giménez et al., 2022). However, a lack of understanding of the design constraints around promoters had made engineering synthetic promoters challenging. Current approaches often require trial and error or high throughput screening to identify functional synthetic promoters (Belcher et al., 2020; Brophy et al., 2022; Brückner et al., 2015; Cai et al., 2020; Moreno-Giménez et al., 2022). A better understanding of the contributions and limitations of core promoters in controlling expression patterns can therefore be essential in engineering better synthetic promoters.

Here, by leveraging publicly available RNA-seq atlases of fifteen angiosperms, we were able to map gene expression pattern onto core promoter type in multiple genomic contexts. While TATA-box-containing promoters are over-represented in conditionally-expressed genes in all of the species we examined, the pattern for Coreless promoters was less clear. In most eudicots, Coreless promoters were over-represented in stably expressed genes, but the opposite trend was observed in monocots. Additionally, by identifying orthologous gene groups within these species, we were able to track changes in core promoter type and expression pattern for groups of evolutionarily related promoters. We found that stably expressed genes are also more likely to have orthologs in other species compared to unstably expressed genes, and the orthologs tend to retain similar expression patterns. Lastly, we show that changes in core promoter types do not explain changes in expression pattern. This evolution-guided approach reveals design rules surrounding core promoter architecture and expression patterns.

## Results

We began this project by identifying species with RNA-seq Atlases, which we defined as datasets containing at least ten different tissue samples and with samples that represented at least two distinct developmental stages. Details regarding the dataset and their references can be found in Supplemental Table S1. Figure1A shows a phylogenetic tree of the fifteen species that fit our criteria, which spans a range of angiosperms including multiple monocots and eudicots. The datasets were processed through a custom pipeline (Figure1B-D). In brief, Kallisto was used for RNA-seq quantification and MultiQC was used to summarize all the outputs up till DESeq2 (Supplemental Data S7) (Bray et al., 2016; Ewels et al., 2016). For each species, normalized counts from each tissue were then converted to stability information using the coefficient of variation (CV) as a metric. In this analysis, lower CV corresponds to more stable expression, meaning comparable expression in all tissues. Higher CV, on the other hand, means less stable and more tissue-specific expression. To facilitate comparison between species, we used percentile rank of CV as the primary metric, which represents the percentage of CVs that are less than or equal to a given value.

**Figure 1.**
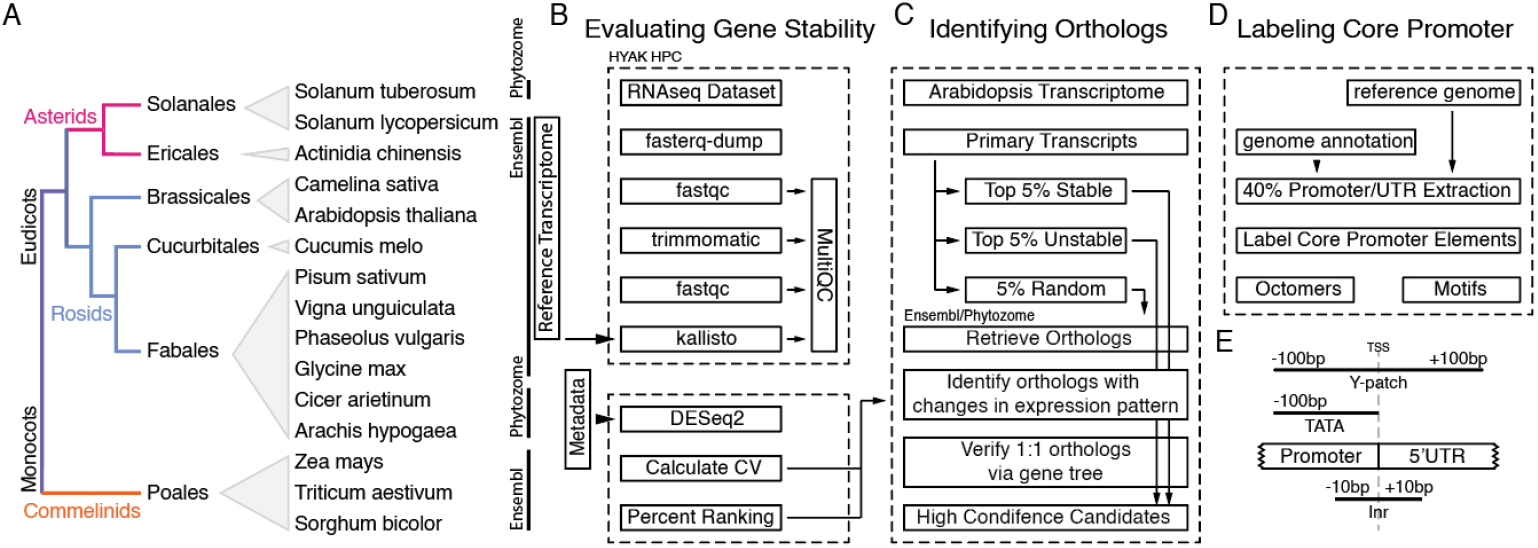
An outline of the bioinformatics pipelines. A) The fifteen angiosperms included in this study and their phylogenetic relationship. B-D) The three major data processing steps performed in the study. Detailed parameters are included in the Methods section. Reference genomes, transcriptomes and gene orthologs were retrieved via either Ensembl (Cunningham et al., 2021) or Phytozome (Goodstein et al., 2012) databases depending on the species. E) Regions searched for each core promoter motif.

To determine whether the characteristic differences in expression patterns between different core promoter types seen in *Arabidopsis* holds across all the species in our dataset, we extracted the - 100bp to +100bp region around the TSS as the “core promoter region” for 40% of all promoters in each species (Figure1D). TATA box, Y patch, and Inr motifs were screened according to methods detailed in Jores et al. 2021. The regions scanned for each motif are more relaxed than their known regions in *Arabidopsis*, as we applied the scan to multiple species and wanted to avoid falsely labeling promoters as Coreless. Illustration of the regions scanned for each core promoter type are illustrated in Figure1E.

Forty percent of all promoters for each species were labeled as either TATA or Y patch. If a promoter did not contain either element, we labeled them as “Coreless”. It is important to note that the definition of Coreless promoters introduced by Yamamoto and colleagues is somewhat more strict than the definition used here, as they also screened for the relatively rare CA and GA core promoter elements (Yamamoto et al., 2009). We then plotted the distribution of CV for each species, broken down by core promoter types (Fig. 2). Similar results for Y patch, Inr and a random set of promoters that serve as a control are in Supplemental Figure S2.

**Figure 2.**
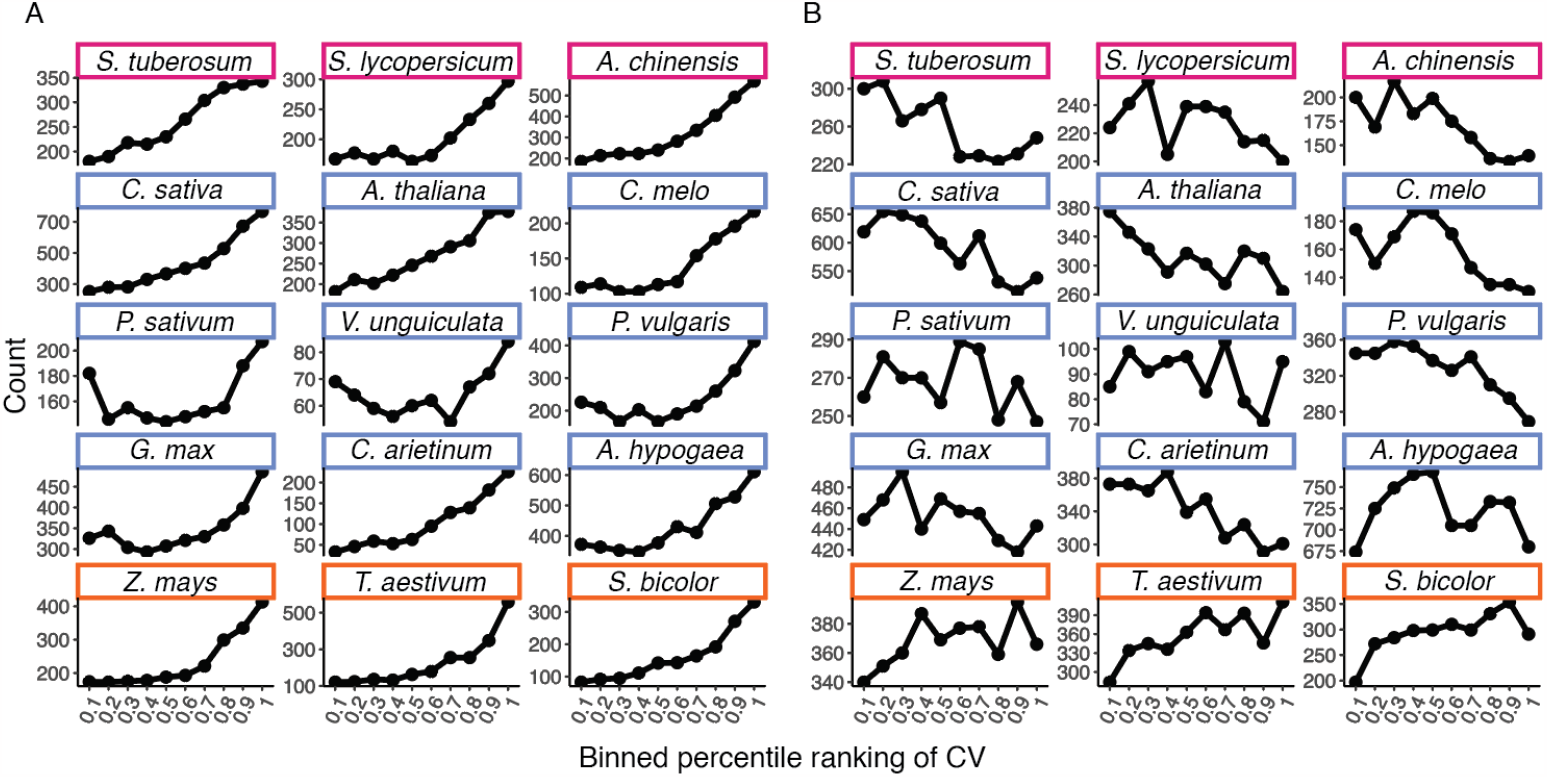
Distribution of relative specificity or uniformity of TATA-box-containing and Coreless promoters. Higher Coefficient of Variation (CV) rankings indicate more specificity, while lower CV rankings indicate more uniformity. A random subsampling of forty percent of promoters from each species are shown here. A) TATA-box containing promoters, and B) Promoters termed Coreless as they lacked both TATA-box and Y-path motifs. Colors correspond to phylogeny shown in Figure 1A.

Using microarray data, Yamamoto and colleagues had found that Coreless promoters are under-represented in genes that responds to stimulus (i.e. more constitutively expressed) (Yamamoto et al., 2011). However, we did not see the same trend until we removed the lowest expressing transcripts from the analysis (transcripts with an average of less than 1 read). These extremely low read counts are likely to be unreliable and an analysis of the weak-expressing genes that we removed revealed that they bias towards higher CV when compared to the rest of the genes in the dataset (Supplemental Figure S3). This same minimum read number requirement was then applied to the rest of the species.

Overall, the expected trend of TATA box-containing promoters being over-represented in unstable genes is observed across all the species analyzed (Fig. 2). In contrast, the trend of Coreless promoters being associated with more stably expressed genes was weaker and only observed in a subset of the eudicots. The monocots (*Zea mays, Triticum aestivum*, and *Sorghum bicolor*) all exhibited a strong trend of Coreless promoters associating with unstable genes (e.g., those with higher CV values), along with an enrichment of Y patch-containing promoters being associated with stable expression (Fig. 2 and Supplemental Figure S2). This inverted pattern could be explained in two ways given that a promoter not labeled as containing a TATA box or Y patch is labeled as Coreless. Under this classification scheme, an apparent enrichment by one category of promoters could reflect a surplus of that type of promoter in a particular CV ranking bin or a depletion of the other two promoter categories in that same bin. The latter explanation seems more likely for the Y patch promoters in monocots, but further experimental tests are required to fully resolve this question. The surprising pattern of Coreless genes “flipping” their behavior in monocots might also reflect an as yet undefined promoter element that is lumped into the Coreless category here. For example, there may be slight differences in TATA motif, as has been described for maize (Mejía-Guerra et al., 2015). Accounting for this known source of variation, we did not see any significant decrease in the Coreless trend towards conditionally-expressed genes (Supplemental Figure S2).

To determine whether core promoter type is tightly linked to expression stability for a given gene, we identified a set of orthologous genes (Figure1C). *Arabidopsis thaliana* is the most well-annotated genome, and it has 47,684 transcripts with a non-zero transcript count in at least one of the sampled tissues. Of this total, we retained only the primary transcripts of each non-mitochondrial and non-chloroplast gene, resulting in a final total of 26,842 genes. The top 5% most stable and top 5% least stable genes were selected based on CV, along with a randomly selected control set of equal size (n=1343 genes in each category). The sets of genes were used to query the Ensembl or Phytozome database for orthologs in the rest of the 14 species in our dataset (Cunningham et al., 2021; Goodstein et al., 2012). The orthologs were searched for in the database where their reference transcriptome was downloaded to ensure matching of the target transcript name with the transcript counts. Orthologs of *Arachis hypogaea, Cicer arietinum, and Solanum tuberosum* were found using Phytozome, and the remaining species were found in Ensembl.

Orthologous genes tended to retain their expression pattern across species (Fig. 3A). While orthologs corresponding to the random set of *Arabidopsis* genes were spread quite uniformly across distribution of CV rankings, the orthologs of the top 5% stable set of *Arabidopsis* genes were skewed heavily towards the more stable, lower percentage CV rankings. The orthologs of the 5% least stable set of *Arabidopsis* genes showed a more subtle skew towards higher CV ranking. This trend was more visible in some species than others, partially due to the overall lower gene counts. One notable trend was that the least stable gene set retrieved significantly fewer orthologs compared to the random or most stable gene sets (Fig. 3B). This is possibly because stable genes are associated with more fundamental cellular functions, and therefore more likely to be conserved across species (Klepikova et al., 2016). Following a similar logic, unstable genes tend to be more tissue-specific, and therefore are more easily lost during species divergence.

**Figure 3.**
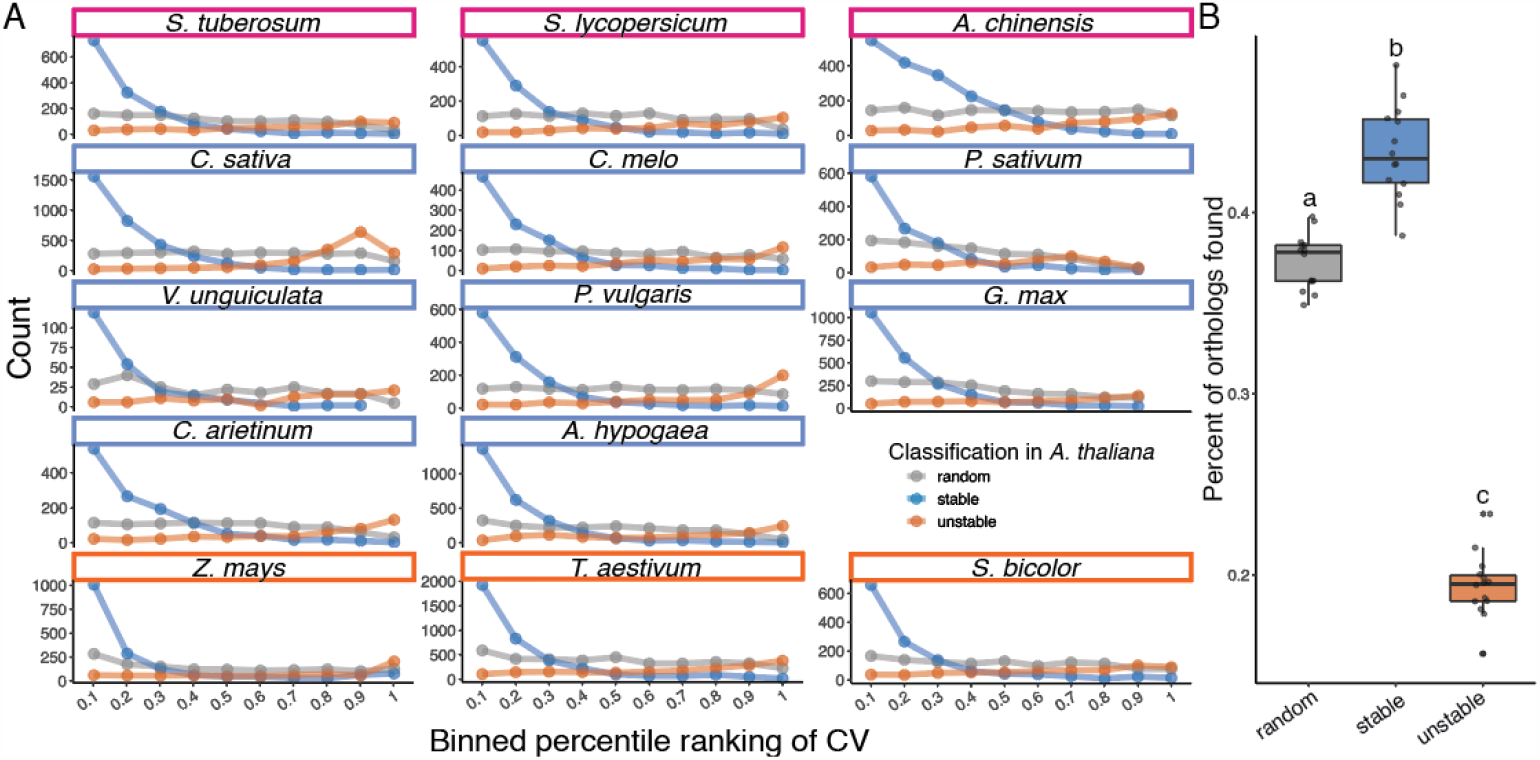
Genes that show uniform expression in *A. thaliana* tend to behave similarly in other species. A) Distribution of CVs for orthologs of stable (blue), unstable (orange) or random (grey) *A. thaliana* genes. The color of boxes around species names corresponds to Figure1A. B) Percent of orthologs found for each set of *A. thaliana* genes for each species. Each dot corresponds to a single species. Statistical tests were performed by one-way ANOVA followed by Tukey HSD. All three groups are significantly different from one another.

Even when looking at genes that fell at the tail ends of the expression stability distribution from *Arabidopsis*, we could find orthologs positioned across the full range of CV rankings (Fig. 3A). In other words, expression stability of a given gene can vary dramatically across species. To investigate this further, we curated a set of evolutionarily-related genes that showed this type of switching behavior. Starting with the set of all the orthologs retrieved through Ensembl and Phytozome, we first filtered the target orthologs to count only the highest expressing transcript for each gene, thereby limiting each gene to a single representative transcript. We filtered the list of orthologs to include *Arabidopsis* transcripts that had only a single ortholog found in the transcriptome of each other species. We considered any target transcripts that crossed the 50th percentile in CV as “changing expression pattern”, and we limited the *Arabidopsis* transcripts to those where transcripts changed expression pattern in at least two different species. These changes were mapped onto the phylogenetic tree to identify clusters where changes could be associated with a specific node.

Gene trees were built for the most promising candidates, and when more than one ortholog was found in the target species, those genes were removed from further analysis (Fig. 1C). These stringent parameters maximize the likelihood that the remaining candidates are true orthologs, and that any changes in expression pattern could be biologically significant. Seven high-confidence orthologous gene groups were found with three *Arabidopsis* transcripts (AT3G17020.1, AT3G18215.1, AT4G40045.1) that are from the top 5% stable genes list and four *Arabidopsis* transcripts (AT1G04700.1, AT5G17400.1, AT5G18910.1, AT5G20410.1) from the top 5% unstable genes list. A summary of the filters and numbers of target orthologs as well as *Arabidopsis* query transcripts left after each step can be found in Supplemental Table S4.

The promoters for these seven sets of orthologs were extracted and TATA, Y patch, Inr motifs were screened for as described above (for clarity, this analysis will be referred to as Motif Scan) (Figure1D). In parallel, these promoters were also screened for TATA, Y patch, Inr, CA, GA octamers as defined in Yamamoto et al. 2009 (Octamer Scan), and an illustration of the regions scanned for each octamers can be found in Supplemental Figure S5. Comparing the two methods, the Motif Scan resulted in more identified core promoters due to its more relaxed parameters. Only two promoters were labeled as Y patch by the Octamer Scan but not the Motif Scan. A core promoter element was considered present if either method returned a positive result (Supplemental Table S6). Within each orthologous gene group, changes in the presence of TATA or Y patch elements did not appear to correlate with changes in expression patterns (Fig. 4). In each group, there are examples of promoters having the same core promoter type but different expression patterns, as well as cases of promoters having the same expression pattern but different core promoter types. Since there were only seven TATA-box-containing promoters (∼15.5% of the promoters), we were not able to observe instances where two related TATA-box containing promoters having different expression patterns, but there are multiple instances where changes in presence of TATA motif did not change expression pattern. This result suggests that the presence or absence of a TATA or Y patch is not sufficient to change expression pattern.

**Figure 4.**
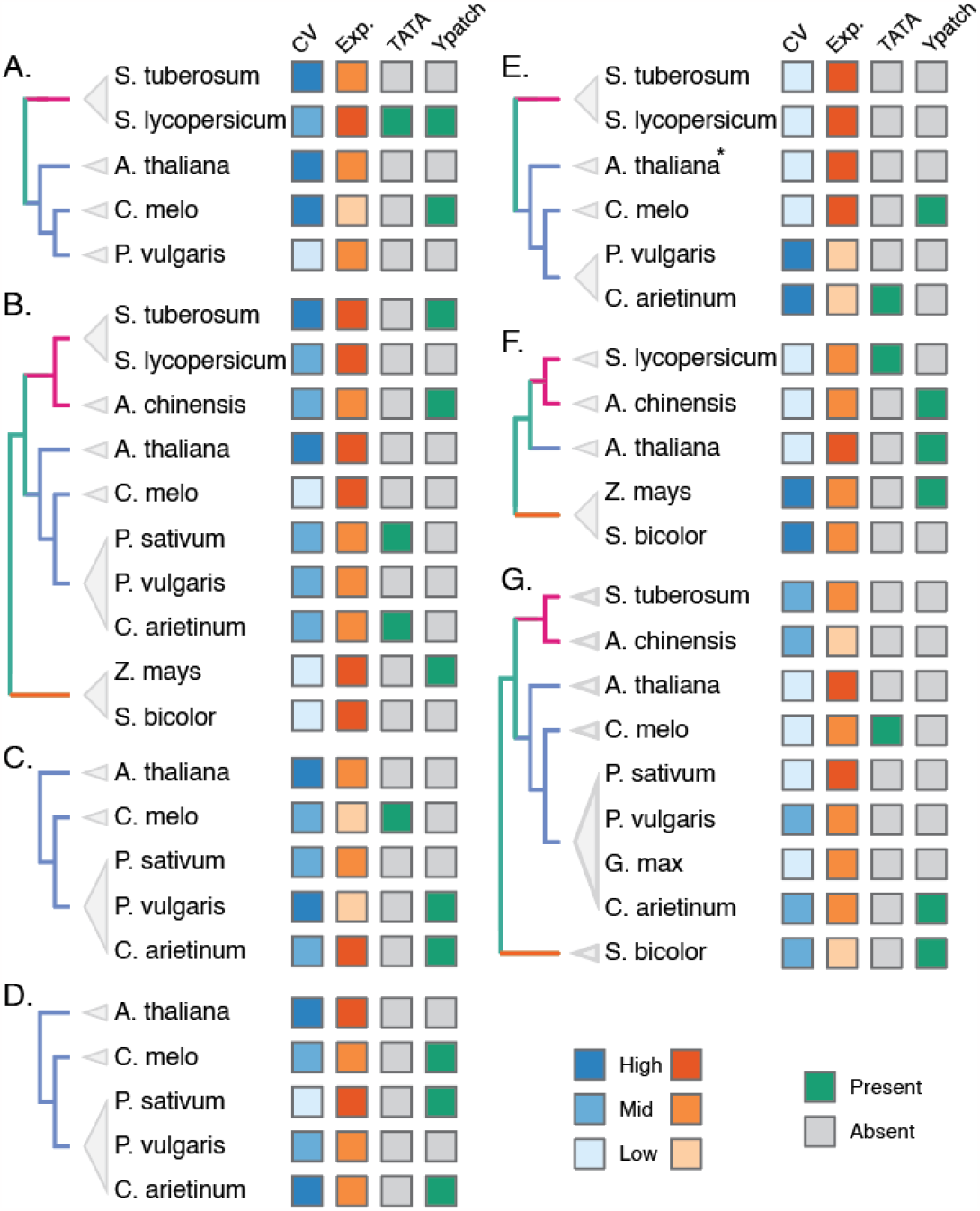
Individual gene trees where expression stability changes can be observed. A-D) The gene is unstably expressed in *A. thaliana* but stably expressed in another species. E-G) The gene is stably expressed in *A. thaliana* but unstably expressed in another species. CV and expression strength (Exp.) is grouped by percentile ranking of 0.66∼1.00 (High), 0.33∼0.66 (Mid), or 0.00∼0.33 (Low) and color coded accordingly. Presence (green) or absence (grey) of TATA and Y patch motifs are indicated. \**A. thaliana* has no identifiable core promoter identified as the intergenic region is only 8 bp.

## Discussion

Understanding the rules that govern the performance of natural promoters could inspire the construction of synthetic promoters that are able to retain their behavior over multiple generations in transgenic plants. Here, we mined RNA-seq atlases from fifteen different angiosperms to extract patterns connected to the relative specificity or uniformity of gene expression across developmental stages and tissue types. We found that the previously observed trend that TATA-box-containing promoters are over-represented in conditionally expressed genes is highly conserved. In contrast, the relative uniformity versus specificity of expression from Coreless promoters is not as well conserved. Coreless promoters from eudicots analyzed in this study were, in general, more highly associated with stable expression patterns. Coreless promoters from monocot species, however, exhibited the opposite trend. In addition, we found that promoters tend to maintain their expression pattern across species, with the caveat that stably expressed genes are more likely to have identifiable orthologs when compared to unstably expressed genes. Lastly, by tracking expression pattern and promoter type within the evolutionary trajectory of individual genes, we could test the hypothesis that promoter architecture is responsible for the level and pattern of gene expression. We found that none of the core promoter types screened for in this work are consistently associated with changes in expression pattern or strength. This suggests that while there may be a correlation between promoter architecture and transcription parameters, the underlying molecular mechanism that determines whether a gene is conditionally or specifically expressed remains unknown.

While the general trend that TATA-box-containing promoters are found in genes that are only expressed in specific times and/or locations was highly conserved, close study of single gene phylogenies reveals that the TATA-box is not the determinant of this expression pattern. The overall lack of pattern for TATA and Y patch motifs on the phylogenetic tree also suggest that the gain and loss of these promoter elements, at least in the genes studied here, are sporadic events that do not experience strong positive selection for maintenance. In the future, it would be interesting to add the additional dimension of tracking the relative conservation versus divergence of the coding regions of the genes associated with each promoter type; however, the small number of promoters in each category would likely limit the potential to detect a clear pattern.

From a synthetic biology perspective, there are two major implications from the analysis described here. First, the hope of finding strong, constitutive natural promoters that work across diverse species may be even more challenging than we originally thought. For example, it is unlikely that there are natural promoter architectures that will work equally well as constitutive promoters in monocot and eudicot crops. Second, and more hopefully, our analysis suggests that the approach currently being taken by multiple labs for engineering synthetic promoters is likely to find solutions that work well across species (Belcher et al., 2020; Brophy et al., 2022; Cai et al., 2020; Moreno-Giménez et al., 2022). The overall scheme of many of these groups is to take a core promoter region containing a TATA-box, and then add natural cis-elements or synthetic transcription factor target sequences. We found that the same core promoter could support widely varied expression patterns. This is consistent with the emerging hypothesis that cis-elements contribute more to expression pattern than the core promoter itself (Cai et al., 2020), and that any desired expression pattern can be achieved regardless of core promoter type. Why Coreless promoters are enriched in constitutively expressed genes in eudicots, and whether this mode of regulation leads to greater robustness of expression pattern over time, will require a more detailed understanding of transcription initiation events at a range of promoters in multiple species.

## Methods

### Phylogenetic tree

A phylogenetic tree was constructed referencing NCBI’s Taxonomy Browser and Li et al. 2021.

### RNA-seq dataset processing

RNA-seq atlases were located in the NCBI Sequence Read Archive (SRA) database. The references for the datasets can be found in Supplemental Table S1. The individual datasets were retrieved using sratoolkit-3.0.1 prefetch followed by fasterq-dump functions. Fastqc-0.11.9 were used to generate a QC report for each dataset. Trimmomatic-0.39 were used for adaptor and low quality ends trimming using the following settings: ‘SLIDINGWINDOW:4:20 MINLEN:36’.

ILLUMINACLIP files TruSEq3-PE-2.fa was supplied for paired end data and TruSEq3-SE.fa were supplied for single end data. Reference transcriptome were downloaded from the Ensembl Plants (http://plants.ensembl.org/index.html) for *Arabidopsis thaliana, Camelina sativa, Cucumis melo, Glycine max, Phaseolus vulgaris, Pisum sativum, Vigna unguiculata, Sorghum bicolor, Zea mays, Solanum lycopersicum, Actinidia chinensis, Triticum aestivum*. and Phytozome (https://phytozome-next.jgi.doe.gov) for *Arachis hypogaea, Cicer arietinum, and Solanum tuberosum* (Cunningham et al., 2021; Goodstein et al., 2012). An index file was generated and the reads aligned and counted using Kallisto-0.44.0 with ‘-o counts -b 500’. For single end data, Fragment Length and Standard Deviation were required, but the information is difficult to locate, and so a default value of ‘-l 200 -s 20’ were used across the board.

Another Fastqc was performed on the trimmed files, and a final MultiQC-1.13 were run on the entire folder encompassing all the log files that Fastqc, Trimmomatic, and Kallisto generated. The MultiQC report was inspected to ensure the trimming step improved read quality and there were no major warnings.

### Normalizing count, Calculating CV and Percent Ranking

#### (Relevant files: 1_Metadata_from_RUNselector.Rmd, 2_MOR_Normalization.Rmd)

Using an R script, the raw counts for each species were normalized using the DESeq2 package using a metadata file curated from the original study for the RNA-seq datasets. The coefficient of variation across all samples for a given atlas was used as a metric for stability for each gene, and the percentile ranking for each gene was calculated. The geometric mean for each gene was also calculated across all samples.

### Extracting intergenic region and 5’UTR

#### (Relevant files: 3_ExtractPromUTR(ALL_Transcripts).ipynb, 8_ExtractPromUTR(Orthologs).ipynb)

Gff3 annotation files and reference genomes were downloaded from Ensembl or Phytozome depending on where the reference transcriptomes were retrieved from. 40% of transcripts were selected from the total transcriptome and their intergenic region and 5’UTR were extracted from the Gff3 annotation. Intergenic region and 5’UTRs of identified orthologs were extracted in a similar manner.

### Labeling core promoter types

#### (Relevant files: 4_Label_Promoters.Rmd, 9_Motif_Scan.Rmd, 10_Octamer_Scan.ipynb)

Motif Scan: Intergenic regions and 5’UTR sequences are trimmed to only regions to be scanned for each core promoter types: TATA box (−100 to TSS), Y patch (−100 to +100), and Inr (−10 to +10). Intergenic regions shorter than 100bps were excluded from analysis. Each regions were scanned for their respective motifs according using motif files as well as methods outlined in (Jores et al., 2021). A motif is considered to be present when the relative motif scores are above 0.85.

Octamer Scan: Intergenic regions and 5’UTR sequences were trimmed based on the positions relative to the TSS outlined in Yamamoto et al. 2009 (TATA, −45 to −18; Y Patch, −50 to +50; CA, −35 to −1; GA, −35 to +75). Each region was scanned for the presence of octamer motifs from the TATA, Y patch, GA, and CA lists outlined in Yamamoto et al. 2009. If the specified region contained at least one motif for a given promoter type, it was labeled as positive.

### Ortholog Analysis

#### (Relevant files: 5_At_gene_ranking.Rmd, 6_Identifying_orthologs.Rmd, 7_Processing_orthologs.Rmd)

The *Arabidopsis* transcriptome was filtered to only include primary transcripts, and mitochondria as well as chloroplast transcripts were removed. Top 5% stable genes by CV, bottom 5% stable genes by CV and a random set of 1343 genes (5%) were randomly selected.

Using biomaRt in R, the Ensembl and Phytozome databases were queried for orthologs for the selected set of *Arabdiopsis* genes for each species (Durinck et al., 2009). Orthologs from *Arachis hypogaea, Cicer arietinum*, and *Solanum tuberosum* were retrieved from Phytozome, and the rest of the species from Ensembl. For analysis in Figure3B, significance test of done by ANOVA followed by Tukey’s HSD. For each target gene that matched to an *Arabidopsis* transcript, only the highest expressing transcript was kept. If an *Arabidopsis* transcript retrieved more than one orthologs from a target species, these pairs of orthologs were removed from analysis. We only kept orthologous gene groups that had a “change” in expression pattern, defined as crossing the 50^th^ percentile CV, in two target species, and the remaining candidates were manually mapped onto the phylogenetic tree to identify gene groups that had changes in expression pattern that are consistent with the tree. This means having changes in expression pattern that are mostly found in the same clade. Gene trees were built for these candidates using blast-align-tree (https://github.com/steinbrennerlab/blast-align-tree) and the candidate lists were further trimmed based on the gene trees to ensure a 1:1 relationship between all members in the gene group.

## Data availability

All scripts and datasets necessary to perform the analysis in the article are available at https://doi.org/10.5061/dryad.9w0vt4bmk

## Acknowledgements

We thank Dr. Alexander Leydon, and Janet Solano Sanchez for careful reading of the manuscript, and Dr. Adam Steinbrenner for advice on identifying orthologs. We also thank other members of the Di Stilio, Imaizumi, Steinbrenner, and Nemhauser lab for their feedback on this project. This work was supported by the National Science Foundation (IOS-1546873), the National Institute of Health (R01-GM107084) and the Howard Hughes Medical Institute Faculty Scholar Award.

## Author Contributions

Experimental design and analysis by EJYY, CJM and JLN. Research performed by EJYY and CJM. Manuscript written by EJYY, CJM and JLN.

